# Like mother, like daughter? Phenotypic plasticity, environmental covariation, and heritability of size in a parthenogenetic wasp

**DOI:** 10.1101/2022.12.02.518902

**Authors:** Alicia Tovar, Scott Monahan, Trevor Mugoya, Adrian Kristan, Walker Welch, Ryan Dettmers, Camila Arce, Theresa Buck, Michele Ruben, Alexander Rothenberg, Roxane Saisho, Ryan Cartmill, Timothy Skaggs, Robert Reyes, MJ Lee, John Obrycki, William Kristan, Arun Sethuraman

## Abstract

*Dinocampus coccinellae* (Hymenoptera:Braconidae, Euphorinae) is a solitary, generalist Braconid parasitoid wasp that reproduces through thelytokous parthenogenesis, an asexual process in which diploid daughters emerge from unfertilized eggs, and parasitizes over fifty diverse species of coccinellid ladybeetles worldwide as hosts. Here we utilized a common garden and reciprocal transplant experiment using parthenogenetic lines of *D. coccinellae* presented with three different host ladybeetle species of varying sizes, across multiple generations to investigate heritability, plasticity, and environmental covariation of body size in D. coccinellae. We expected positively correlated parent-offspring parasitoid regressions, indicative of heritable size variation, from unilineal (parent and offspring reared on same host species) lines, since these restrict environmental variation in phenotypes. In contrast, because multilineal (parent and offspring reared on different host species) lines would induce phenotypic plasticity of clones reared in varying environments, we expected negatively correlated parent-offspring parasitoid regressions. Our results indicate (1) little heritable variation in body size, (2) strong independence of offspring size on the host environment, (3) a consistent signal of size-host tradeoff wherein small mothers produced larger offspring, and vice versa, independent of host environment. We then model the evolution of size and host-shifting under a constrained fecundity advantage model of Cope’s Law using a Hidden Markov Model, showing that *D. coccinellae* likely has considerable fitness advantage to maintain phenotypic plasticity in body size despite parthenogenetic reproduction.

## Introduction

Size of an organism is a complex and often plastic trait that is correlated with key adaptive traits such as reproductive success (Bosch and Vicens 2005, Berger et al., 2012), fecundity (Honek 1993), response to varying environments and hosts (Chown and Gaston 2010), developmental rates (Davidowitz et al., 2003), survival (Callier and Nijhout 2013), and greater depredation success (Oliveira et al., 2019). At the same time, larger bodied organisms face challenges such as increased resource need, and strong evolutionary constraints on reproductive tradeoffs (Blanckenhorn 2000, Shine 1988), which set “thresholds’’ on size. Theory therefore predicts that a fecundity advantage for body size only occurs in the presence of energy availability (Shine 1988). The evolution of organismal size has been studied extensively over speciation timescales (reviewed in Hone and Benton 2005), often pointing to multiple independent transitions to larger body size (termed as Cope’s Rule) across diverse animal taxa, indicating that there is no one definitive “pathway” or evolutionary strategy for size among species. Several lines of evidence instead support that plasticity of body size evolves at microevolutionary scales (Maurer et al., 1992), with standing genetic variation providing the basis for adaptability of body size plasticity (Gotanda et al., 2015). In insects, for instance, standing genetic variation determines the range of body size that can be expressed in adults, and the varying environmental conditions during larval development can modify how body size is expressed in adults (Honek, 1993; Schneider et al., 2011). The body size of female insects is correlated with fecundity, and plasticity in body size expression can be an adaptive strategy for some insects (Honek, 1993). Endoparasitoids, however, can only exploit the limited resources defined by their host morphology throughout all larval instar stages (Du et al., 2021). Plasticity in the expression of body size is therefore expected to be a beneficial strategy for endoparasitoids, as it enables each larva to successfully develop in a larger range of host body sizes (Du et al., 2021). P ositive correlations between body size of host and adult offspring are well documented in solitary parasitoid wasps, as parasitoid offspring reared on larger species and sizes of host develop larger parasitoid offspring than their parent (Mackauer and Chau, 2001; Arakawa et al., 2004; Wang et al., 2008).

As tradeoffs are ever present across phenotypes, the plasticity of parasitoid body size occurs as a tradeoff between accessing wider ranges of hosts for generalist parasitoids, but at the expense of producing varying sizes of adult parasitoid offspring (Henry et al., 2006). Parthenogenetic wasps therefore provide an ideal natural experimental system to test hypotheses of plasticity of body size, considering their clonal mode of reproduction that maintains genetic variation, specifically utilizing a combination of common-garden and reciprocal transplant experiments to control for genetic and environmental variation.

The parasitoid wasp, *Dinocampus coccinellae* (Hymenoptera: Braconidae), is a generalist that is capable of successfully parasitizing over fifty species of predatory ladybeetles (Coleoptera: Coccinellidae, subfamily Coccinellinae) across a global distribution (Balduf, 1926; Ceryngier et al., 2018, Fei et al 2023). *D. coccinellae* primarily displays solitary behavior, and is only known to asexually reproduce through thelytoky, a mode of parthenogenesis in which females emerge from unfertilized eggs; with males rarely observed in this species (Slobodchikoff and Daly, 1971; Wright, 1979; Heimpel and De Boer, 2008; Ceryngier et al., 2018). Briefly, thelytoky is a parthenogenetic mode of reproduction in which diploid female adults develop from unfertilized egg clones (Beukeboom et al., 2007; Heimpel and Jetske, 2008; Slobodchikoff and Daly, 1971). There are genetic forms of thelytoky in which no crossing over occurs (apomictic thelytoky or premeiotic doubling) or where the fusion of sister or non-sister recombinant chromosomes form diploid eggs (automictic thelytoky) (Heimpel and Jetske, 2008), regardless restricting genomic variation from parent to offspring.

Characteristic to the Euphorinae subfamily of Hymenoptera, a parasitoid larva of *D. coccinellae* consumes the adipose tissue of a parasitized adult ladybeetle as a koinobiont endoparasitoid, (Balduf, 1926; Orr et al., 1992; Ceryngier et al., 2018), although it has been documented to oviposit within host ladybeetle larvae and pupae (Obrycki et al., 1985). Across the diverse range of host ladybeetles, *D. coccinellae* has been reported to preferentially oviposit in coccinellids which are more mobile, larger, adult, female hosts (Davis *et al*., 2006; Obrycki, 1989). Once an adult *D. coccinellae* locates a sufficient adult ladybeetle, they arch their stinger under the beetle and thrust into the abdomen of the host, injecting clonal daughter egg(s) along with accompanying venom enzymes and the RNA-virus, the *Dinocampus coccinellae* Paralysis Virus (DcPV) (Balduf, 1926; Orr *et al*., 1992; Dheilly *et al*., 2015). This is yet another unique facet of the *D. Coccinellae*, as their life cycle involves an endosymbiotic relationship established with DcPV, an RNA virus in the Iflaviridae family (Dheilly et al., 2015). In concert with host behavior modifications mediated by this virus, *D. coccinellae* then use their captive adult host as a bodyguard to the advantage of the next generation. After approximately a week following oviposition within a host beetle, the larva emerges from its egg into the fat body of the host’s abdomen, where it undergoes four larval instar stages of development (Balduf, 1926). Multiple eggs may be deposited within the same host, which is referred to as superparasitism, which has been documented in several field studies (summarized by Ceryngier et al. 2012). When this occurs, the first larva to emerge crushes the others with its mandibles (Balduf, 1926). In these cases of superparasitism, the survivor then cannibalizes its host-mate(s) as its first meal; otherwise, the larva feeds on adipose tissue and ovaries of coccinellid host throughout development (15-20 days) (Balduf, 1926). Tetratocytes, which originate from the parasitoid’s egg, aid in providing an initial food source, in addition to the host itself (Okuda *et al*., 1995). Following pupation in an external cocoon, the daughter wasp emerges as an adult with fully developed eggs, with some of these females leaving a varying percentage of their hosts alive (Orr et al., 1992). The intricate behavioral relationship between an adult *D. coccinellae* wasp and its host ladybeetle have been described, with successful parasitization, measured as the percentage of emerged daughter wasps as a proxy for fecundity, varying between different host species (Orr et al., 1992). However, little is known about fitness of the emergent parthenogenetic daughter wasps.

A previous study by Vansant et al. (2019), found positive relationships between host and emergent daughter *D. coccinellae* morphology (e.g. dry mass, wing length, abdominal length), which has been found to be the case in a variety of parasitoid wasps and their hosts (Brandl and Vidal, 1987; Mackauer and Chau, 2001; Harvey et al., 2006; Henry et al., 2006; Symonds and Elgar, 2013). The developmental environmental conditions, including resources that a developing parasitoid can uptake from its host, substantially determines the body size phenotype of the emerging parasitoid. Additive genetic effects on the body size phenotypes are expected to be heritable, and the relative importance of the development environment and additive genetic effects in determining the body size phenotypes of *D. coccinellae* are not known. However, as *D. coccinellae* reproduces via thelytoky with little to no genetic recombination, this brings into question the balance between heritability or phenotypic plasticity of body size as a proxy for individual fitness.

Endoparasites can only consume limited nutrient resources from a single host throughout all stages of their development. As *D.coccinellae* feeds on the same adipose tissue resource across many host coccinellids, and female reproductive organs when available, the body size plasticity in *D. coccinellae* reflects the volume of host adipose tissue available for consumption, with larger hosts providing more resources to consume and support the growth of the developing parasitoid. Endoparasitoids therefore benefit from plasticity in body size, as it allows larvae to develop successfully in a larger range of host body sizes (Du et al., 2021). Host non-specificity of the parthenogenetic wasp *D. coccinellae* make this species a good candidate to examine the narrow sense heritability of body size plasticity in an asexual, clonal wasp species across generations and host species.

Given that total phenotypic variation of a trait is composed of genetic and environmental variation, and genetic variation in *D. coccinellae* offspring is limited due to thelytokous parthenogenetic reproduction, we designed two models of reciprocal transplant experiments: in the first, we developed a *D. coccinellae* wasp lineage of at least 4 generations on a single species of host coccinellid and referred to this as a ‘unilineal’ setup; in our second model, we developed a *D. coccinelae* wasp lineage of at least 2 generations on multiple host coccinellid species, reciprocally alternating between three species of host coccinellid of varying size and referred to this as a ‘multilineal’ setup. In a unilineal setup, we hypothesized that adult *D. coccinellae* wasps would have minimal variation in size from environmental effects, as the same host species would keep the environment that each *D. coccinellae* larva develops under relatively constant, in terms of size of host coccinellid and therefore available resources for the developing parasitoid larva to consume during development stages. We expect that restricting the environmental variability to a single host species would produce no relationship between parent and offspring body size, if size is entirely phenotypically plastic, whereas a positive relationship if body size in *D. coccinellae* daughter offspring would be largely due to additive genetic variation. Alternatively, in a multilineal setup, we hypothesize that adult *D. coccinellae* would have relatively more variation in size from the changing host environment for their larvae to develop under. We therefore expect negative relationships between parent and offspring body size if the body size of adult *D. coccinellae* is entirely phenotypically plastic. Additive genetic effects may cause a residual positive correlation between mother and daughter pairs after host size has been accounted for in multilineal lines. The relative importance of additive genetic effects and developmental environment could then be assessed by combining unilineal and multilineal lines in a single analysis.

## Materials and Methods

### Experimental setup

*D*. *coccinellae* wasps used to start the lineages for reciprocal host-transplants were obtained from field collections in Kentucky of parasitized adult *Coccinella septempunctata* (*C. septempunctata* or C7 *-* JJO personal comm.) and from *Hippodamia convergens* (*H. convergens* or H. con) from an insectary in San Marcos, CA. Parthenogenetic lines of *D. coccinellae* were then maintained for at least 4 generations on laboratory populations of three species of lady beetles – C7, H. con, and *Coleomegilla maculata* (*C. maculata* or C. mac) which were obtained from field sites in Kentucky (JJO personal comm.). These beetle populations were maintained on an *ad libitum* diet of *Acyrthosiphon pisum* (pea aphids), which in turn were maintained on fava bean plants (*Vicia faba*) in insect tents in the California State University San Marcos (CSUSM) greenhouse in San Marcos, CA until March 2020. Following the COVID-19 outbreak, all insect tents, subsequent crosses, and experimentation were performed (socially distanced and masked) in AT’s garage in Oceanside, CA. Despite temporary relocation of the experimental setup, all experimental conditions were maintained constant to minimize random effects, including daily variations in temperature and diurnal cycles. The unlineal and multilineal experiments were conducted in a single location, with the same species of three hosts, and therefore approximately evenly distributed across treatments.

In each experimental setup, one adult *D. coccinellae* wasp (‘mother’) was placed into a paper soup cup along with four individual ladybeetle hosts, moth (*Ephestia*) eggs for hosts to feed on and a honey-water soaked cotton ball for both the wasp and beetles to drink from; only one wasp was introduced per each experiential cup setup and was sealed using a mesh sheet and an open-face lid. After the mother wasp oviposited into her hosts and died (which always occurred before her daughters egressed from their cocoons), she was paired with the host that she emerged from for imaging. The remaining four host ladybeetles were fed and tended to until the initial appearance of the cocoon spun by the larval *D. coccinellae* parasitoid. Once finished developing in her cocoon, an adult *D. coccinellae* ‘daughter’ egressed from her cocoon. This ‘daughter’ *D. coccinellae* was then placed in another experimental cup setup as the next ‘mother’ *D. coccinellae* with another four individual ladybeetles of the next type of host coccinellid species, *Ephestia* (moth) eggs for hosts to feed on and honey-water for both host and wasp to drink. In every introduction cup, the life history data recorded were: wasp introduction date, parent removal and collection date, cocoon date (if noticed), daughter eclosion date, and host mortality rate. Mothers were picked at random to oviposit in unilineal or multilineal conditions.

Parasitized beetles were reared until *D. coccinellae* larva egression from the infected host as a cocoon woven between the host legs (Vansant et al., 2019). 92 wasp-host pairs and 40 mother-daughter pairs were collected for morphological observations. An expanded polystyrene foam stage and ruler (mm) was assembled to standardize and scale the photographed parasitoid-host pairs. Using a Nikon dissection microscope, adult *D. coccinellae* wasps were photographed in the lateral position, and the corresponding ladybeetle host was photographed from the dorsal, lateral, and ventral positions. These images were uploaded into ImageJ (NIH) to obtain the following morphometric measurements in mm for the wasp: head length, head depth, thorax length, thorax depth, abdomen length and wing length (Figure 1); and for the host beetle: dorsal body length and depth; lateral body depth, elytron chord length and pronotum length; and ventral pronotum width, and abdominal length and width (Figure 2); based on body segments measured in Vansant et al., 2019. Morphometric measurements were repeated independently by four individuals and averaged to control for observational bias. Each parent and offspring wasps were then paired for regression analysis, in addition to pairing host beetle and emergent wasp measurements.

**Figure 1.**
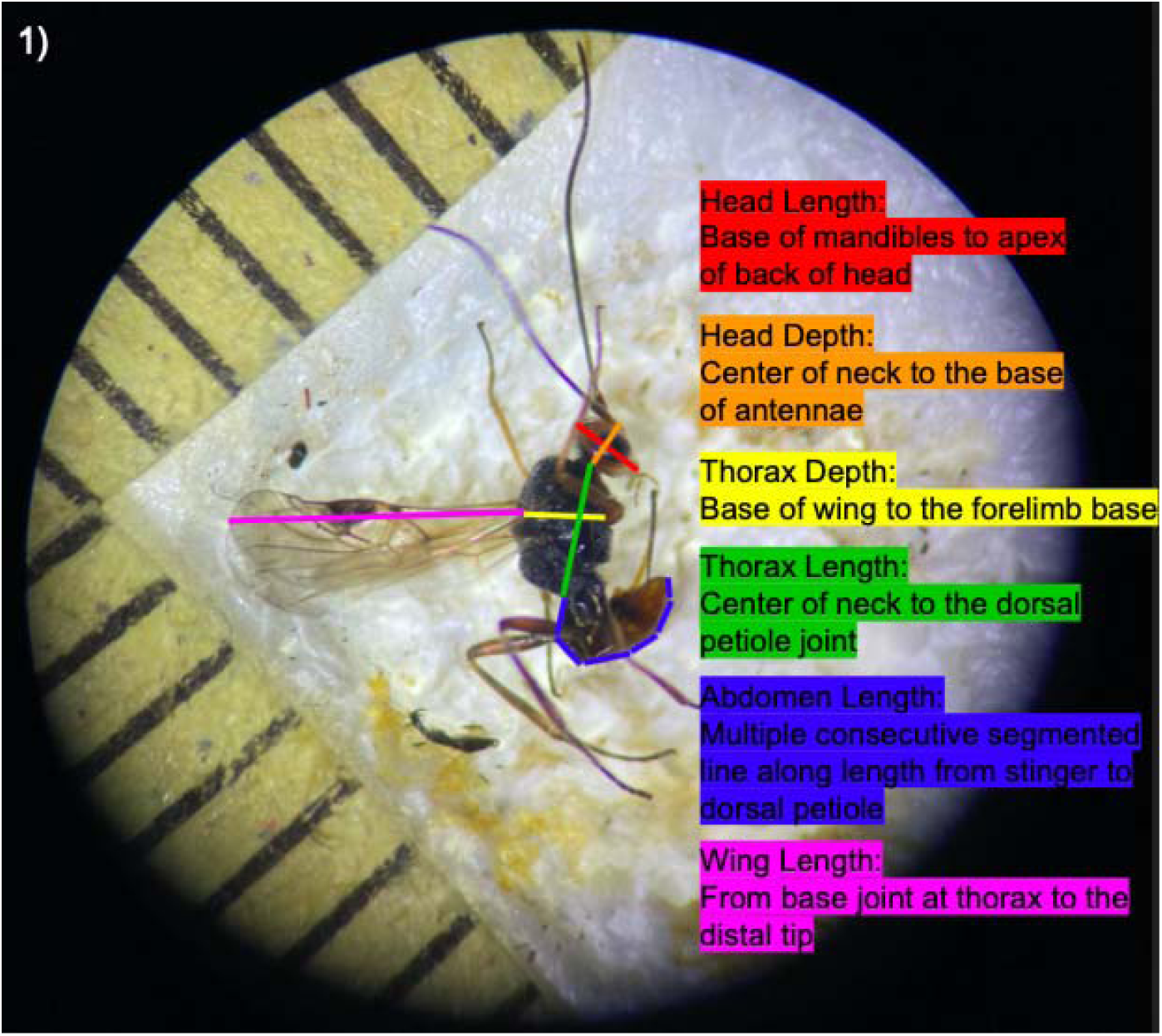
Body segment morphometric traits measured from adult *D. coccinellae* parasitoid wasps, shown in lateral view with a millimeter scale on the stage. These traits were selected based on the morphometric segments outlined in Vansant *et al*., 2019.

**Figure 2.**
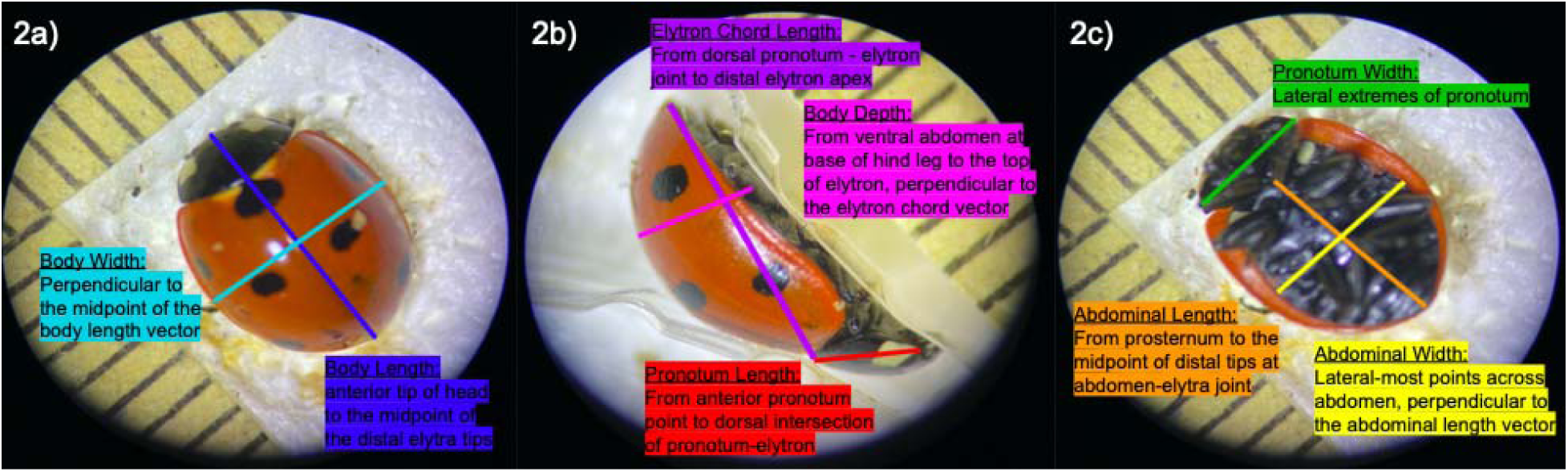
Body segment morphometric traits measured from host lady beetles which the parasitoid *D. coccinellae* egressed from. Shown from dorsal (Fig. 2a), lateral (Fig. 2b), and ventral (Fig. 2c) perspectives, with a millimeter scale on the stage. These traits were selected based on the morphometric segments outlined in Vansant *et al*., 2019. **2a)** “Body Width: Perpendicular to the midpoint of the body length vector”, “Body Length: anterior tip of head to the midpoint of the distal elytra tips.” 2b) “Elytron Chord Length: from dorsal pronotum – elytron join to distal elytron apex,” “Body Depth: From ventral abdomen at base of hind leg to the top of elytron, perpendicular to the elytron chord vector”, and “Pronotum Length: From anterior pronotum point to dorsal intersection of pronotum-elytron.” 2c) “pronotum Width: Lateral extremes of pronotum,” “Abdominal Length: From prosternum to the midpoint of distal tips at abdomen-elytra joint”, and “Abdominal Width: Lateral-most points across abdomen, perpendicular to the abdominal length vector.”

### Statistical analysis

All statistical analyses and visualization were performed in R v.4.2.1 (R core Team), with additional libraries noted below.

Size distributions of all wasp and host ladybeetle morphological variables were visualized as box plots, grouped by the host species from which the wasp eclosed. Differences between host species morphologies were tested using one-way ANOVA followed by Tukey HSD post-hoc procedures for each host morphological variable. Additionally, due to the similarity in sizes of *C. maculata* and *H. convergens* hosts, we grouped them into a “Small” group and compared them to the “Large” *C. septempunctata*. Similarly, box plots, and one-way ANOVA were performed across *D. coccinellae* morphological measurements, with host size as a factor. The multivariate relationship between host morphology and morphology of *D. coccinellae* that developed in the host was assessed using canonical correlation analysis (CCA). The body segment measurements for *D. coccinellae* and their hosts were used (Figure 2). Statistical significance was assessed using an F approximation of Wilks’ Lambda (Rao’s F), using the CCP package in R (CCP version 1.2, Menzel 2022).

Narrow sense heritabilities of wasp morphological variables were measured with parent/offspring regression. Crosses were assigned to either the unilineal group (when parent and offspring host species were the same) or the multilineal group (when parent and offspring host species differed), and type of cross was included as a factor in the parent/offspring regressions. Models were fitted using sum contrasts, such that the slope for the main effect of parent morphology represented the overall parent/offspring relationship across the two different types of cross. The interaction of cross type with parent host morphology tested for differences in slope of parent/offspring relationships between unilineal and multilineal crosses. When a significant interaction occurred, indicating that unilineal and multilineal slopes differed from one another, we used the *emmeans* package (Lenth 2022) to test the slopes for difference from zero. A second set of models with parent host species replacing cross type as a factor in the models, used a similar approach, but differences in parent/offspring slope between the three host species were tested using post-hoc procedures. Parent morphology was not orthogonal to the categorical predictor in either sets of linear models. To determine whether the confounded variation between morphology and the grouping variable (i.e. cross type or parent host species) was responsible for significant relationships between parent and offspring morphology, two alternative models were fitted for each morphological variable with parent morphology entered either before or after the factor, and then the two alternative orders of entry were tested with Type I (sequential sums of squares) ANOVA. Cases in which a significant parent/offspring regression became non-significant after accounting for the factor were noted (Supplemental Data File).

To better understand the association between mother and daughter morphology, Canonical Correlation Analysis (CCA) was conducted relating mother and daughter morphological variables (mass was not included because of missing values, which would have further reduced the sample size). Initially, the canonical correlation between mother and daughter morphology was assessed without accounting for host type. Then, to determine how much of the canonical correlation between mother and daughter was due to either mother or daughter host environment, partial CCA was conducted using the residuals from a MANOVA that included mother’s host, daughter’s host, or the combination of both mother’s and daughter’s host as predictors. Partial CCA that controlled for mother’s host in mother’s morphology and daughter’s host in daughter’s morphology, and for mother’s host in mother’s morphology and combinations of mother’s and daughter’s hosts in daughter’s morphology were also used. Large reductions in canonical correlation between mother and daughter when host was accounted for would indicate that the correlation was principally due to host-mediated effects (e.g. developmental environment, mother’s investment decisions at oviposition), whereas stable patterns of correlation after host effects had been accounted for would be consistent with factors driven by the mother’s state, independent of the host she developed in or oviposited on. All partial CCA analyses were conducted with the *yacca* package (version 1.4-2, Butts 2022), which included a measure of statistical redundancy between the mother’s and daughter’s morphological variables. Statistical significance of canonical correlations was assessed using an F approximation of Wilks’ Lambda (Rao’s F), using the CCP package in R (CCP version 1.2, Menzel 2022).

We additionally employed a Redundancy Analysis (RDA) to gauge the effects of maternal morphometrics and beetle host species on offspring wasp morphometrics (Okasanen et al. 2022). Before analysis we scaled and centered parent and offspring morphometrics using the “scale()” function in base R (R Core Team, 2021). After standardizing our morphometric data, we added parent and offspring beetle host species as categorical environmental predictor variables to the morphology matrix. We constructed a full RDA model with offspring wasp morphometrics as response variables and parent wasp morphometrics, all parent and offspring host species as predictor variables. Subsequently we used the *step* function from the stats package in R to evaluate all combinations of reduced sets of predictors to obtain ones that explain the most variation in the response variables. The reduced set of predictors were then assessed for collinearity with the *vif.cca* function that is part of the *vegan* package (Oksanen et al., 2022). The terms with a score of less than 20 were chosen to build a reduced RDA model. We evaluated the global, axis and term significance of the RDA model with the *anova.cca* function with additional parameters “permutations=9999”, “by=”axis”” and “by=”terms”” respectively. Additionally we obtained the unbiased adjusted R squared value using the *RsquareAdj* function to determine the amount of variance in the response matrix explained by the predictor matrix. Subsequently, we constructed an ordination plot to visualize parent morphology and offspring host species predictors in relation to offspring response variables using the *ordiplot* function from the *vegan* package (Oksanen et al., 2022).

### Modeling host-shifting as an evolutionary strategy - is size heritable?

In order to further explore the idea that host-shifting in parthenogenetic parasitoid wasps might be an evolutionary strategy, we designed a model that describes maintenance of phenotypic plasticity in size under a constrained fecundity variation of Cope’s Law. Under this Hidden Markov Model, fitness is optimized as a function of (a) the efficacy of parasitization and oviposition, (b) number of viable offspring, (c) size of the offspring, (d) size of the host, and (e) availability of the host. Host size was modeled as a binomial distribution, with “Small” and “Large” size thresholds, while parasitoid size was varied continuously, with thresholds. Transition probabilities between the hidden states were simulated based on a cost to host-shifting, such that it occurs only while optimizing a Gaussian fitness landscape (thresholded such that fitness can never be 0, wherein the wasp population would go extinct) that uses the parameters described above. The evolution of size was modeled under Cope’s Law as a selection gradient parameter, beta, such that beta = 0 indicates no selection on size, positive beta indicating selection for larger size. All simulation parameters are described in Table yyy. We then simulated 100 generations under both scenarios, to understand (1) the frequency of host-shifting as a function of fitness, (2) heritability of size under a constrained fecundity model. The maximum *a posteriori* host states were then estimated using a Viterbi algorithm using the “GaussianHMM” function in *numpy*.

### Does offspring fitness change depending on the heritability of size?

We also performed a separate set of simulations for 1000 generations under neutrality (beta = 0) and under positive selective constraint (beta = 0.15, which is the median linear selection gradient across animalia - Kingsolver and Pfennig 2004) to assess variation in offspring fitness versus narrow sense heritability (h^2^ = 0.1, 0.3, 0.9) to understand support for Cope’s Law under host abundance variation.

## Results

### Experimental results

To establish that the three separate ladybeetle species do indeed provide parasitoid *D. coccinellae* larva with significantly different environmental conditions to develop within, we generated box plots and ran one-way ANOVA between the ladybeetle species across each body segment (Figure 3a-i). *C. maculata* and *H. convergens* were much more similar in size than either were to *C. septempunctata*, and some morphological variables were not significantly different between them (body length, abdominal length, mass). But, once *C. maculata* and *H. convergens* were grouped together under the ‘Small’ label and *C. septempunctata* being the ‘Large’ label (Figure 3a-i), all differences were statistically significant.

**Figure 3a-3h.**
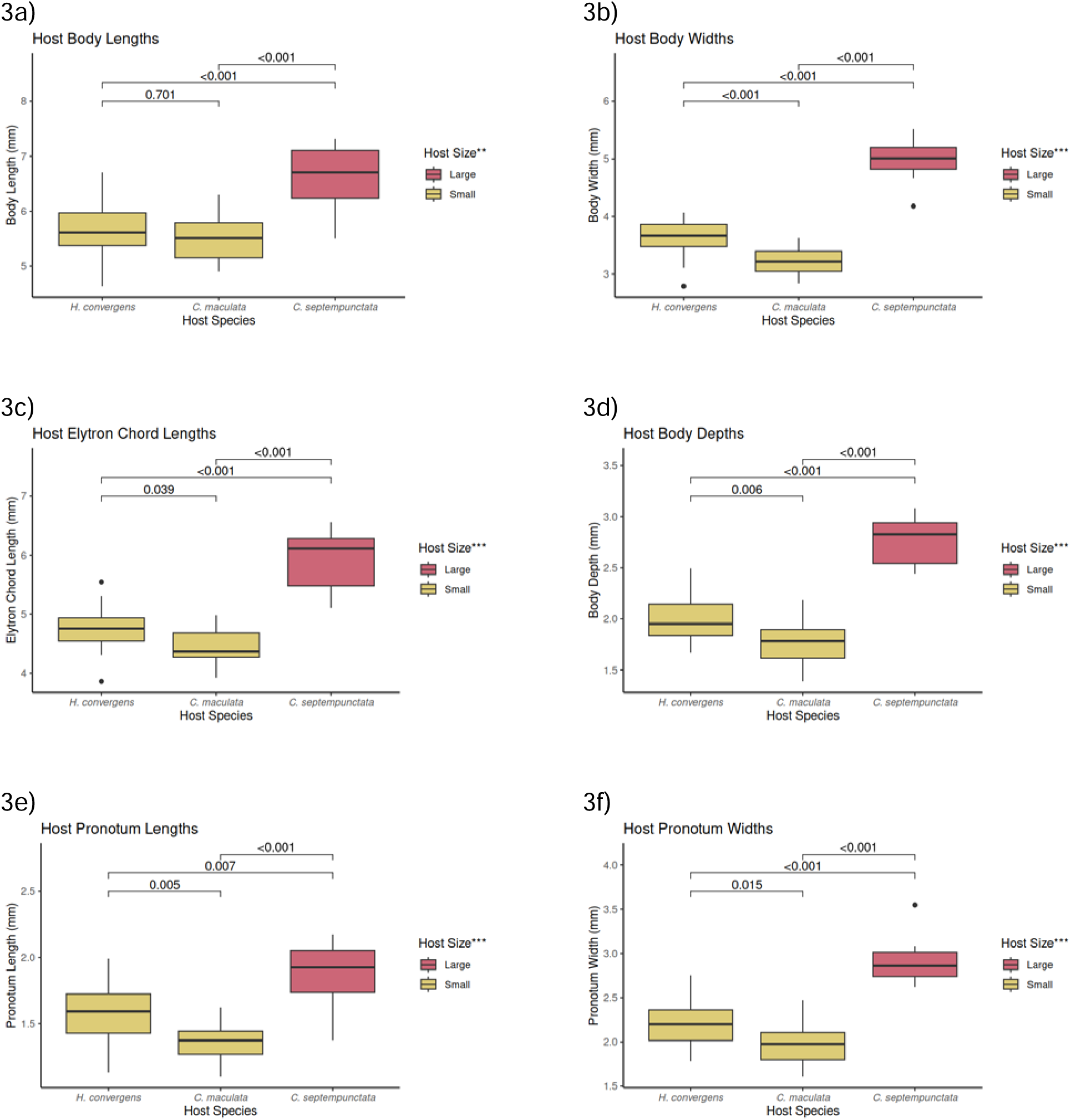
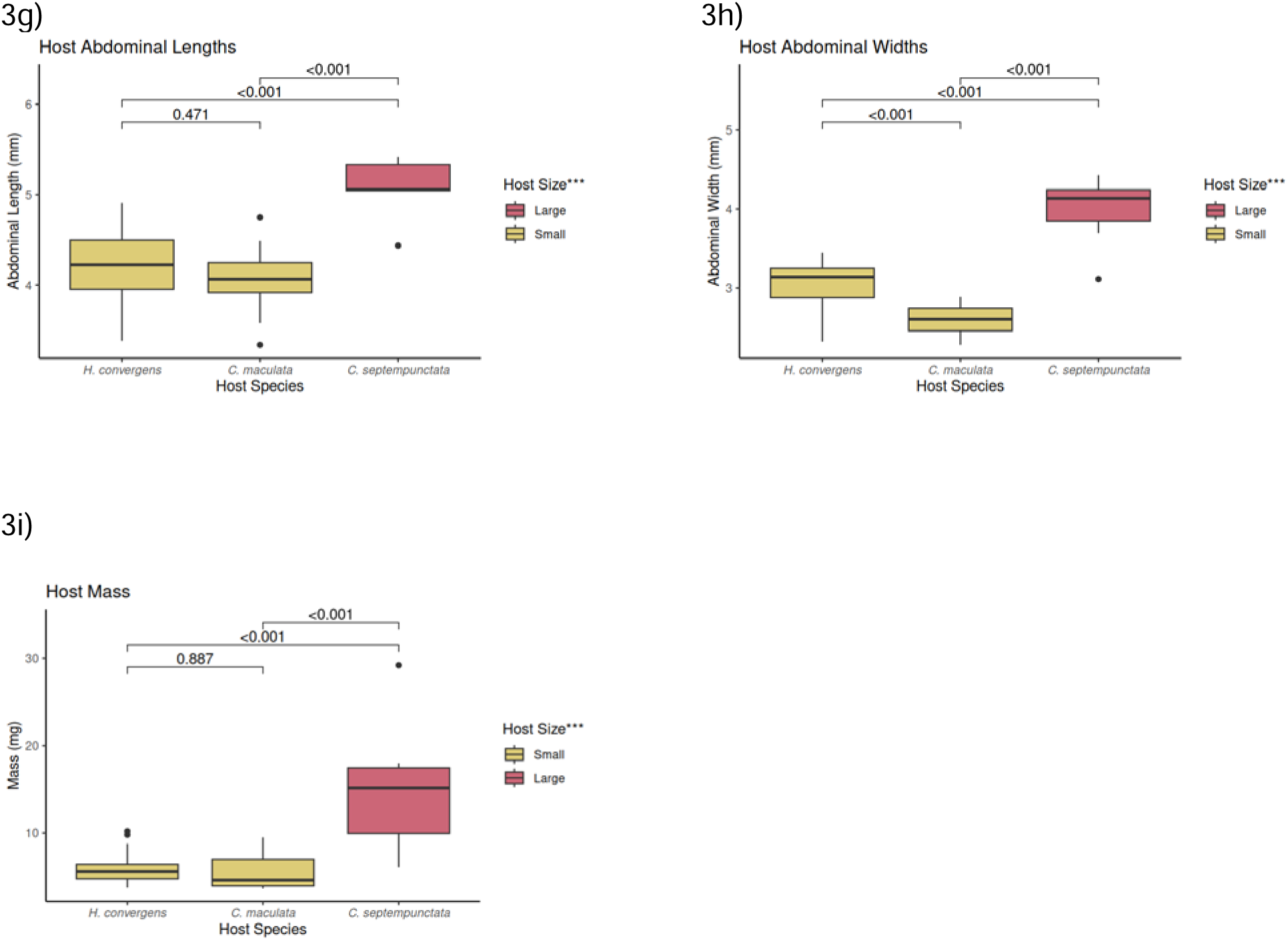
Boxplots of morphometric variables measured for *Coleomegilla maculata*, *Hippodamia convergens*, and *Coccinella septempunctata* host ladybeetles across the dorsal, lateral, and ventral image viewpoints. P-values from Tukey post-hoc comparisons are indicated above box plots. Significance level for comparisons between small (combined *H. convergens* and *C. maculata*) and large (*C. septempunctata*) ladybeetles are indicated as asterisks on legend title (* < 0.05, ** < 0.01, *** < 0.001).

*D. coccinellae* grown on different host sizes only differ in abdominal length, (Figure 4). The two strongest correlations between host and *D. coccinellae* morphological variables are between ladybeetle abdominal width and *D. coccinellae* head depth (r = 0.51), and between ladybeetle abdominal width (V) and *D. coccinellae* thorax length (r = 0.45). The multivariate correlation between host and *D. coccinellae* morphology is much higher; the first canonical correlation coefficient R = 0.83 was statistically significant (p = 0.014), but the second through sixth were not (p > 0.05). The first canonical correlation axis represents a positive relationship of host abdominal width (loading = −0.94) with *D. coccinellae* size across every variable but head length (loadings ranging from −0.21 for thorax depth to −0.56 for head depth; Figure 5).

**Figure 4.**
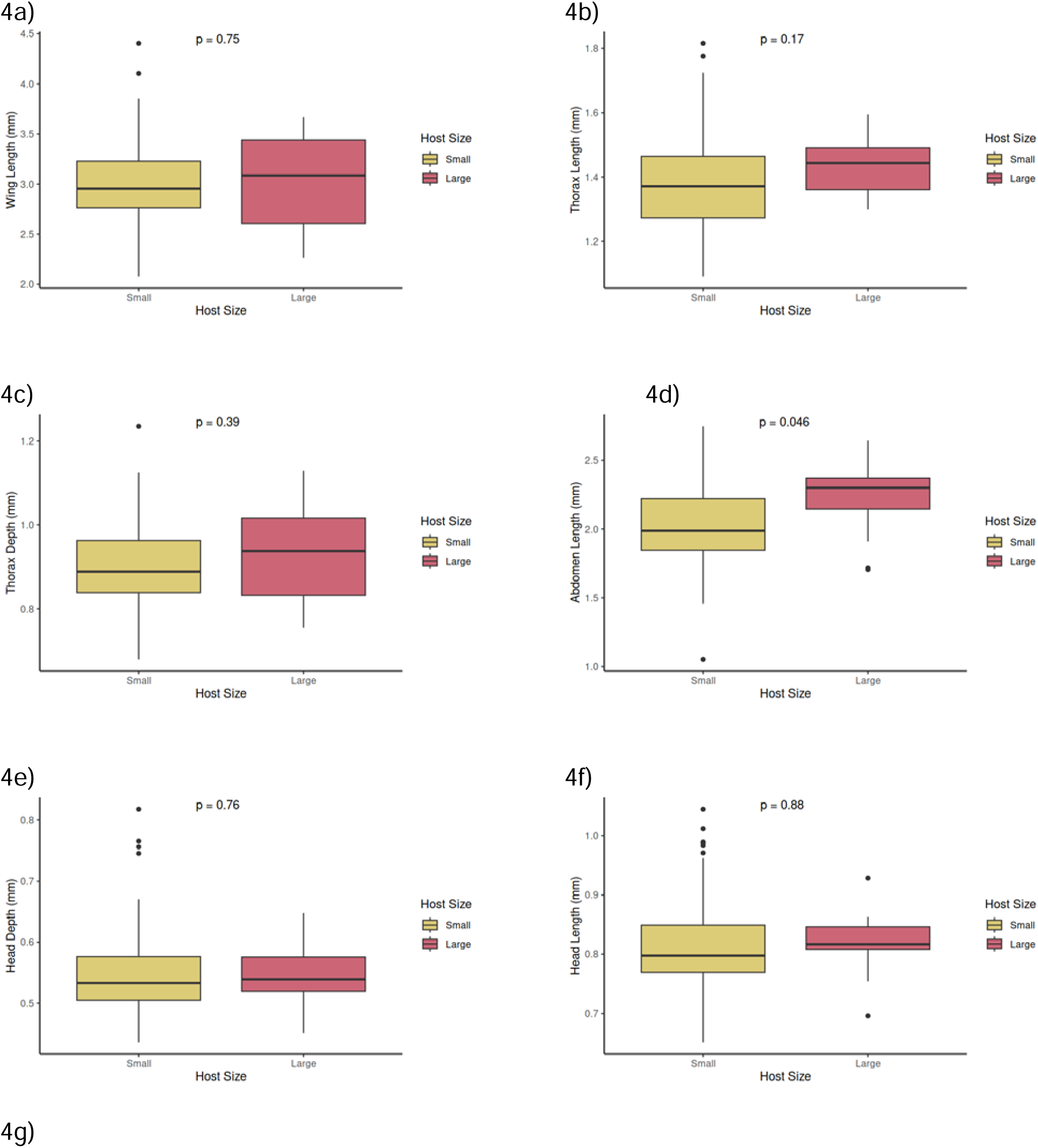
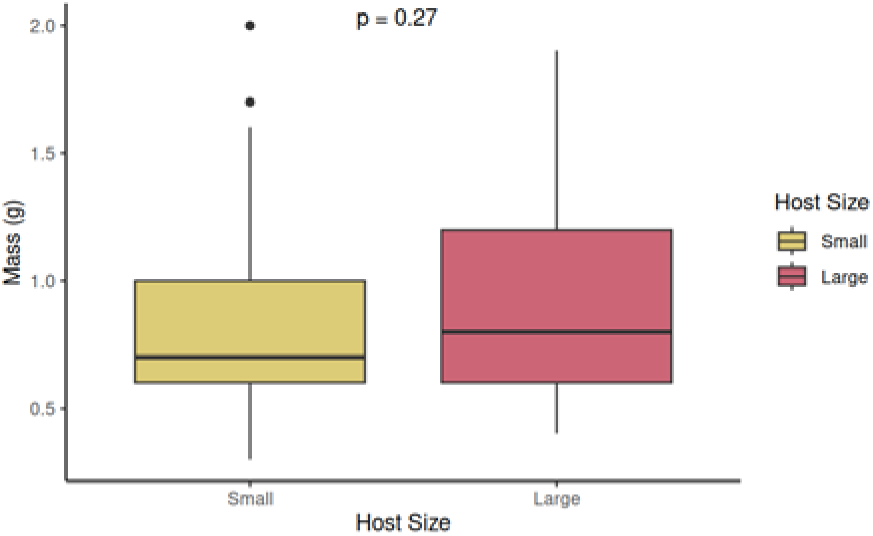
Boxplots for morphometric variables measured for *D. coccinellae*, separated by which host ladybeetle species the parasitoid egressed from (‘Small’ and ‘Large’ host categories are the same for the host ladybeetle analyses). The p-values are comparisons of means by one-way ANOVA.

**Figure 5.**
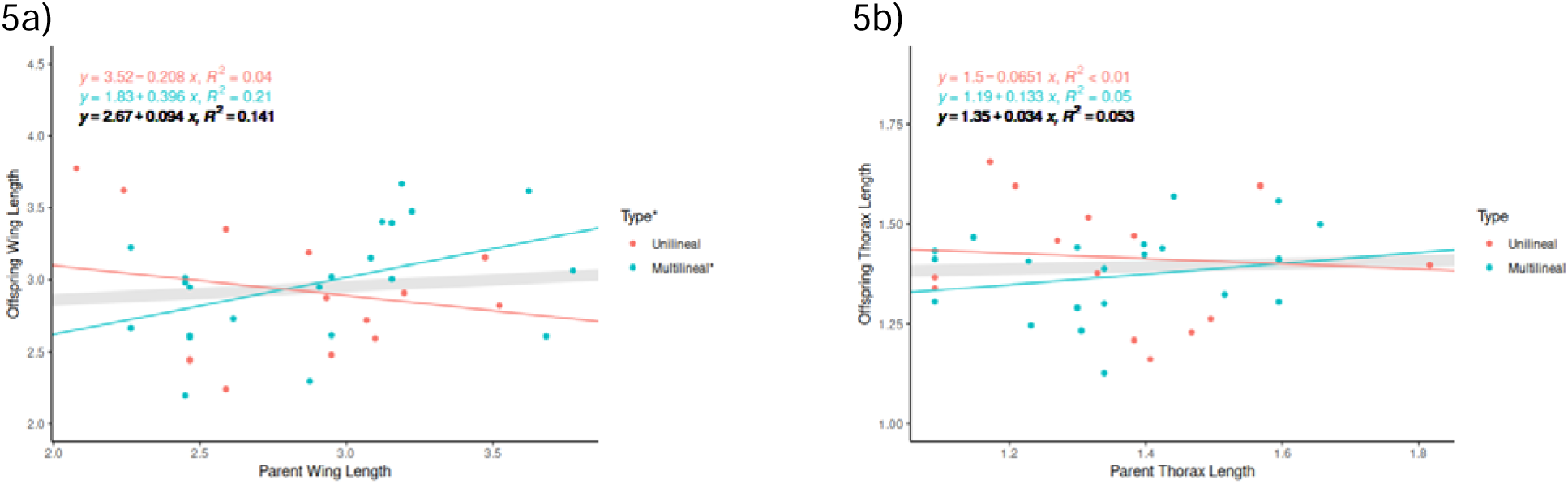
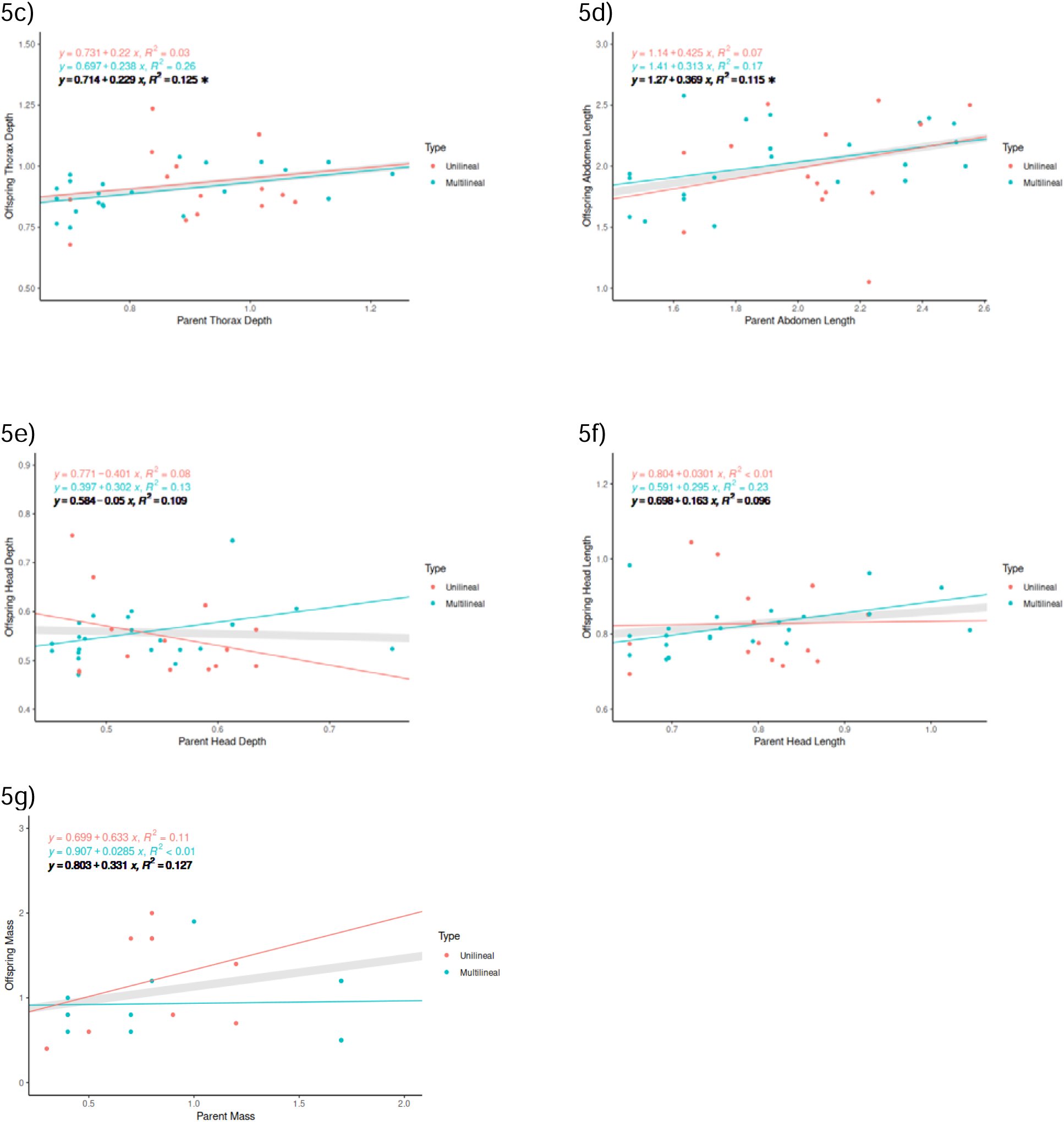
Analysis of covariance relating offspring morphology to parent morphology, grouped by type of cross. Unilineal crosses used the same host species for both parent and offspring, and multilineal crosses used different host species for parent and offspring.. The gray line gives the overall relationship between parent and offspring, and its regression equation and coefficient of determination are shown in black (asterisks indicate p < 0.05). Unilineal and multilineal regression equations and coefficients of determination match the color for their lines and plot symbols (an asterisk on the legend title indicates a significant Type x parent morphology interaction, and an asterisk next to a type label indicates a slope significantly different from zero for that group).

Narrow sense heritability (h^2^) across each body segment measurement was captured by the slope of the line of the main effect of the parent morphological variable parent-offspring linear models (Figures 6a-g and 7a-f). Additionally, the parent-offspring regressions in Figures 6a through 6g distinguish between unilineal and multilineal crosses, whose slopes are represented by the parent morphology by cross type interaction. Four out of seven unilineal parent-offspring regressions indicate a slight positive slope, but for some variables they are negative (wing length, thorax length, head depth), and none of the unilineal slopes are significantly different from 0 (n = 15). Alternately, regression slopes for all multilineal parent-offspring pairs uniformly display a slight positive relationship [total n = 25], and in the one variable (wing length) that had a significant difference in slopes between cross types only the positive multilineal slope significantly differed from zero (Fig. 5a). The overall parent/offspring relationship, combined across cross type, was significant for thorax depth (Fig. 5c) and abdomen length (Fig. 5d), but the slopes did not differ by type of cross for these two variables.

**Figure 6.**
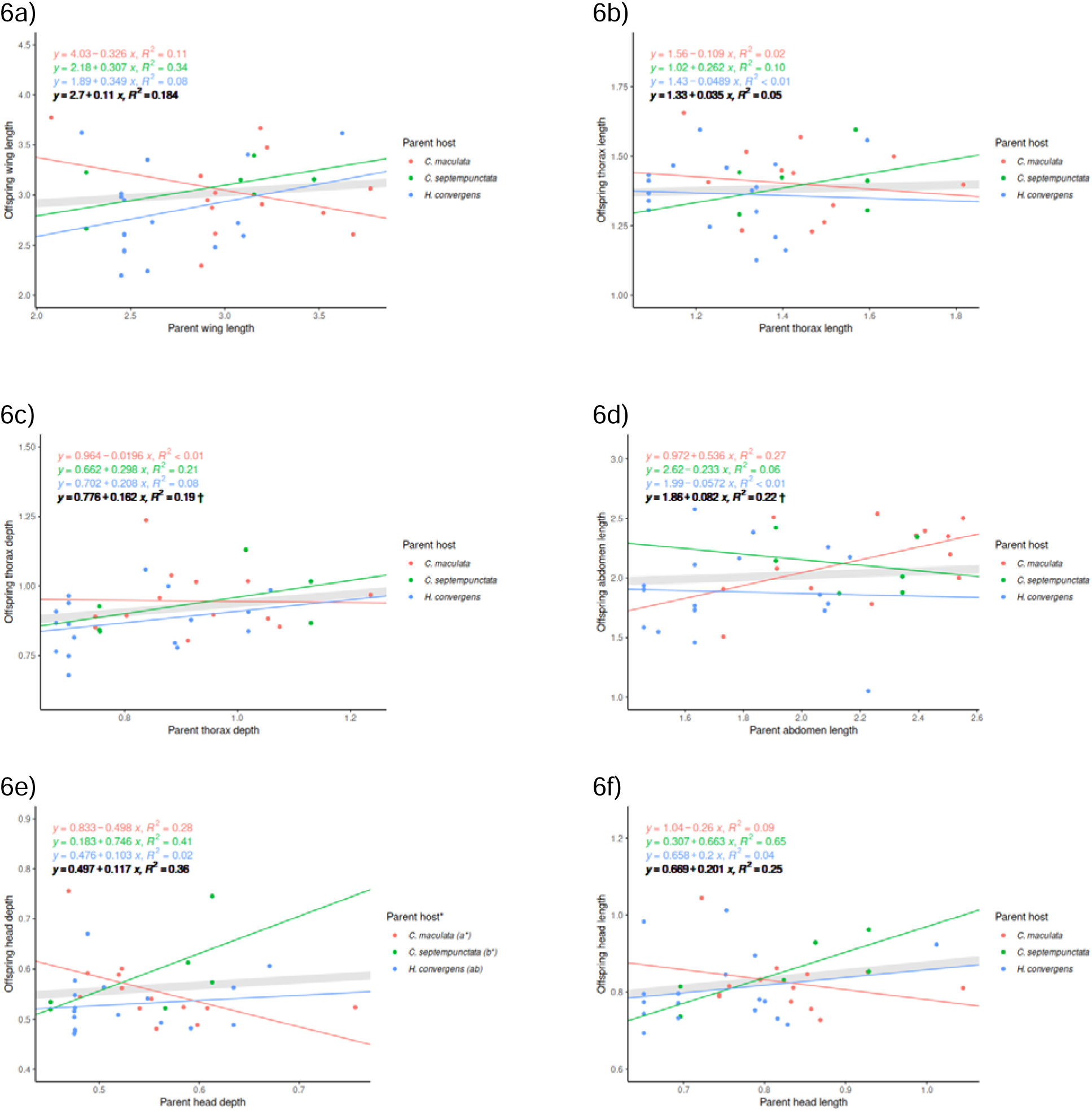
Parent offspring analysis of covariance models, grouped by parent host. The overall relationship between parent and offspring is shown with a thick gray line, and its regression equation and coefficient of variation are shown in black. Regression equations and coefficients of variation are colored to match lines and plot symbols for host species. Symbols indicate: * = p < 0.05, † = p < 0.05 only when entered first using sequential sums of squares. Letters after host species names in 7e are post-hoc comparison letters, and * indicates slopes that are significantly different from 0.

**Figure 7.**
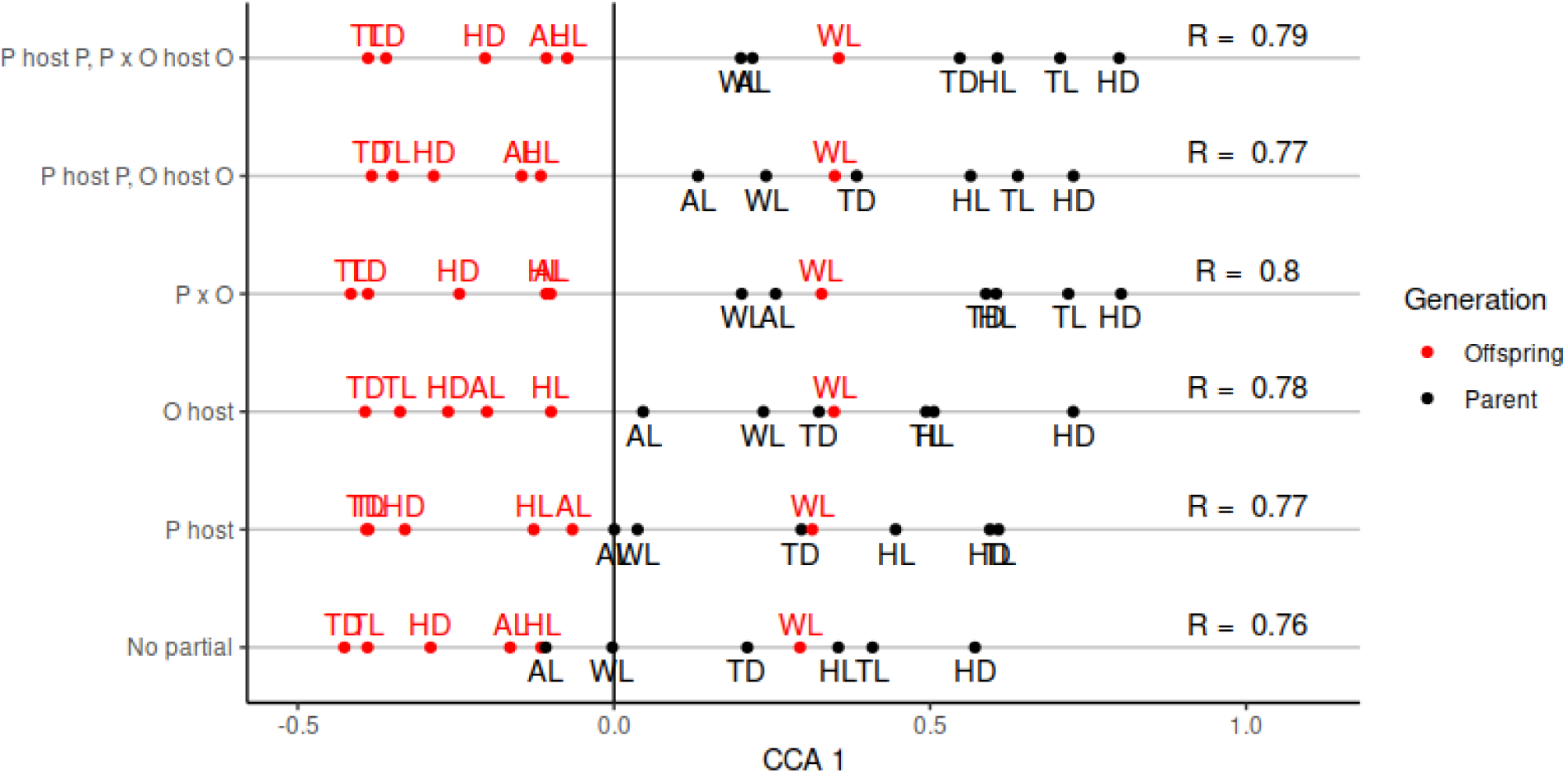
Summary of the results of canonical correlation analysis between mother and daughter morphology. Each model is shown as a row, labeled along the y-axis. Points give variable loadings (correlation between variable and CCA1 axis), and are labeled by variable abbreviation. The canonical correlation coefficient for CCA1 is given to the right of the points.

Using parent host species to group the data yielded significant parent/offspring relationships only for thorax depth (Fig. 6c) and abdomen length (Fig. 6d) as before, but confounding between parent host species and parent morphology made these relationships dependent on the order that parent morphology was entered into the model; when they were entered first they were significant, but when they were entered second they were not. Parent hosts only differed in their slopes for head depth (Fig. 6e), with *C. maculata* and *C. septempunctata* differing from one another, and neither differing from *H. convergens*. The negative slope for *C. maculata* was significantly different from zero, as was the positive slope for *C. septempunctata*.

For every CCA of parent and offspring morphology only the first canonical axis was statistically significant (Rao’s F approximation, p-values ranging from 0.002 to 0.02). All of the models yielded a qualitatively consistent pattern (Figure 7), in which the parent loadings were primarily (for the “No partial” model that did not account for any host effects) or entirely positive and offspring loadings were negative except for wing length. The canonical correlation coefficients were also very consistent, ranging from a low of 0.76 for the “No partial” to a high of 0.8 for the model in which the combinations of mother and daughter host were accounted for in both the mother and daughter morphologies. The contrast between mother and daughter loadings increased in any model that the mother’s or daughter’s host was accounted for, with all of the mother’s loadings becoming positive for those models.

The offspring, parent wasp morphology and parent and offspring wasp vs beetle host relationship was examined by Redundancy Analysis RDA. Testing the global significance of the full RDA model with all parent morphometrics, parent and offspring species as predictors yielded a non-significant result (F = 1.6271, p = 0.0685). The stepwise reduction of predictors from the full RDA model and subsequent evaluation of reduced models by AIC yielded a model with head depth, thorax depth and offspring with an adjusted R squared value (R^2^=0.158). The global RDA significance of the reduced model was significant (F = 2.8353, p = 0.009). The first RDA axis was significant (F = 8.5682, p= 0.026), however, the other RDA axes were insignificant at a FPR cutoff of 0.05 (p > 0.05). Offspring host and parent thorax depth were both determined to be significant terms in the RDA model. The results of the ordination indicate that offspring wasps raised in a unilineal setup tended to have smaller thorax depth measurements compared to their parents. In contrast, offspring wasps raised in multilineal setups tended to have larger thorax depths. Additionally, given the direction of the predictor vectors, the model ordination shows a positive correlation between offspring head depth and thorax depth and parent head depth and thorax depth (Fig. 8).

**Figure 8:**
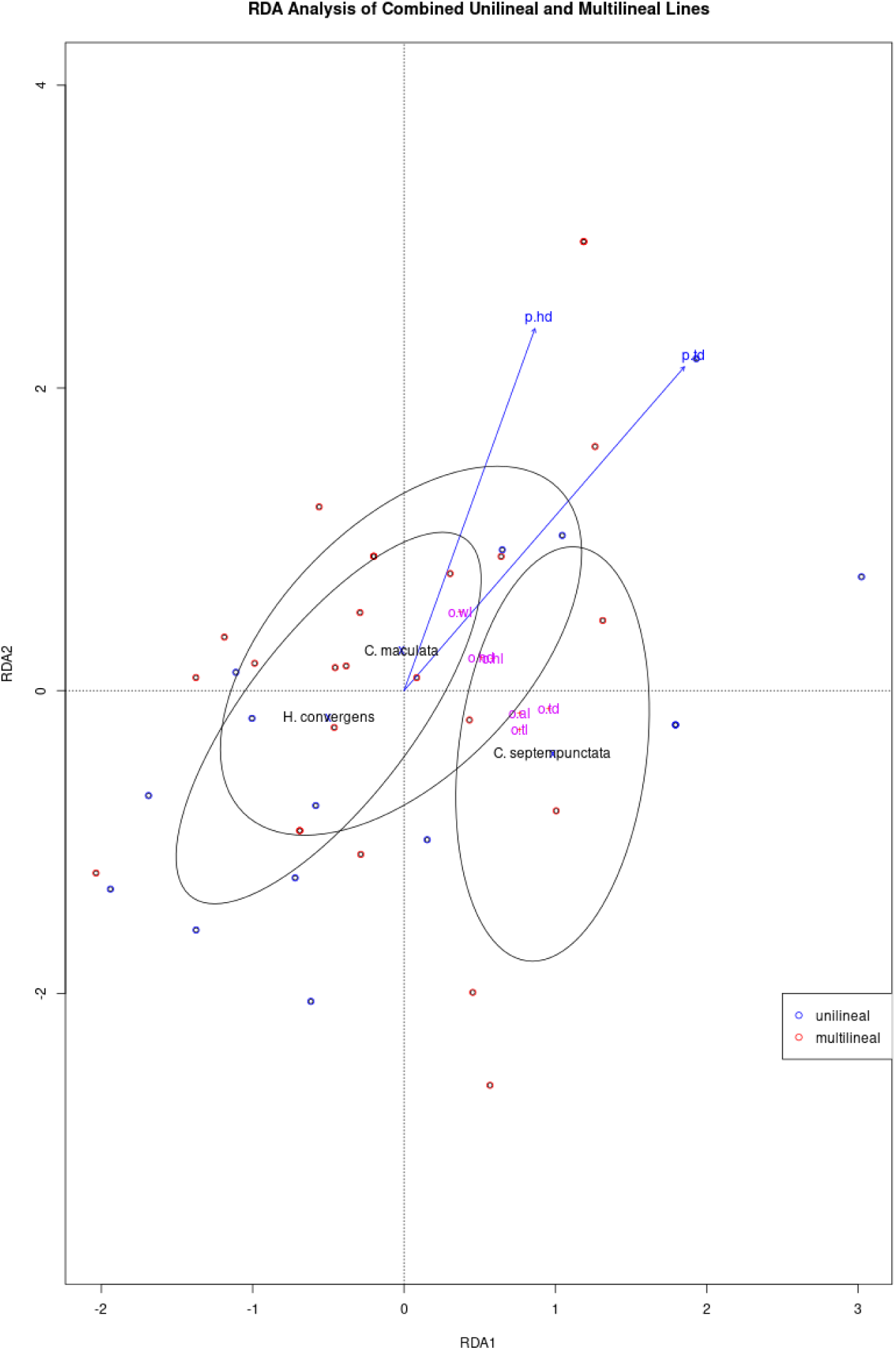
Redundancy analysis (RDA) with reduced set of predictors of the response of *D. coccinellae* offspring wasp morphology to multilineal or unilineal growth environment. Blue and red circles represent individual offspring wasp samples recorded under unilineal or multilineal setup respectively. Parent head depth and thorax depth represented by blue vectors ‘p.hd’ and ‘p.td’ respectively. Offspring morphology response variables represented in pink by ‘o.wl’ = Wing length, ‘o.hl’ = Head length, ‘o.hd’ = Head depth, ‘o.al’ = Abdomen length, ‘o.td’ = Thorax depth and ‘o.tl’ = Thorax length. Offspring host categorical predictors represented as ellipsoids.

### Simulation results

HMM simulations of wasp body size evolution as a function of (a) host shifting strategies, (b) host availability, (c) parasitization efficiency, (d) parasitization success, and (e) offspring fitness indicate that our HMM accurately predicts host-shifting of wasps in conjunction with host-size (Fig. 9(A)), even when host availability predicts a transition of host-state (from large to small or vice versa). Correspondingly, efficacy of parasitization and offspring fitness are correlated with (Fig. 9(B)), and vary as a function of host abundance. For instance, greater abundance of hosts around generation 76 also correlates with greater parasitization efficacy, and therefore greater offspring fitness. Body size is also predicted to be highly heritable (Fig. 9(C) narrow sense heritability h^2^ = 0.88) but less fit (offspring fitness < 0.2), under a model of Cope’s Law with constrained fecundity (selection gradient β = 0.15, with selection favoring larger host sizes). To investigate the relationship between offspring fitness and the heritability of size, repeating simulations with varying narrow-sense heritabilities of body size (h^2^ = 0.1, 0.3, and 0.9) under neutrality (β = 0.0) and fecundity constraint on host size (β = 0.15) clearly indicate that offspring body size however is predicted to have fitness advantages under plasticity, rather than under a model of high heritability (Fig. 10(A) and 10(B)).

**Figure 9:**
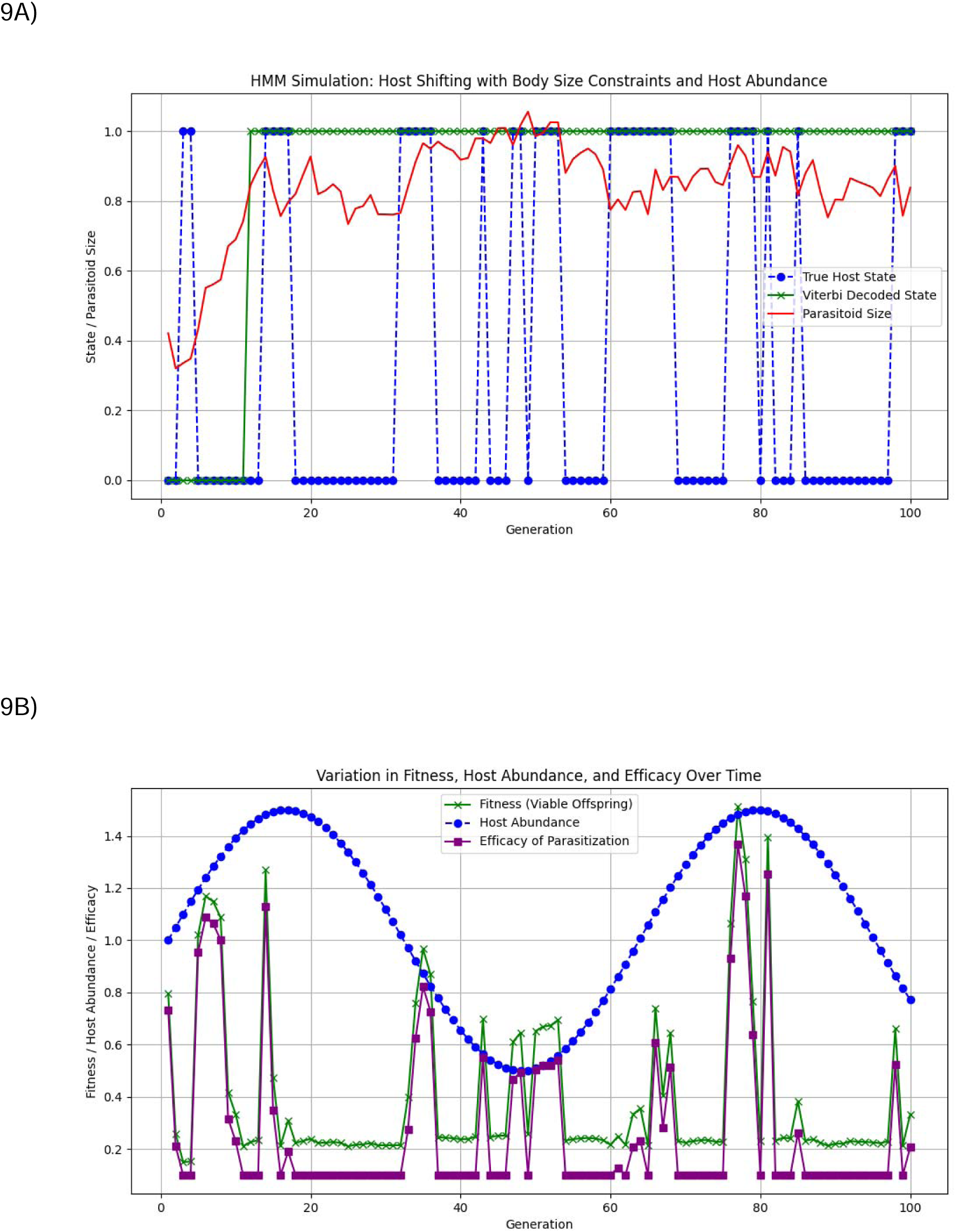
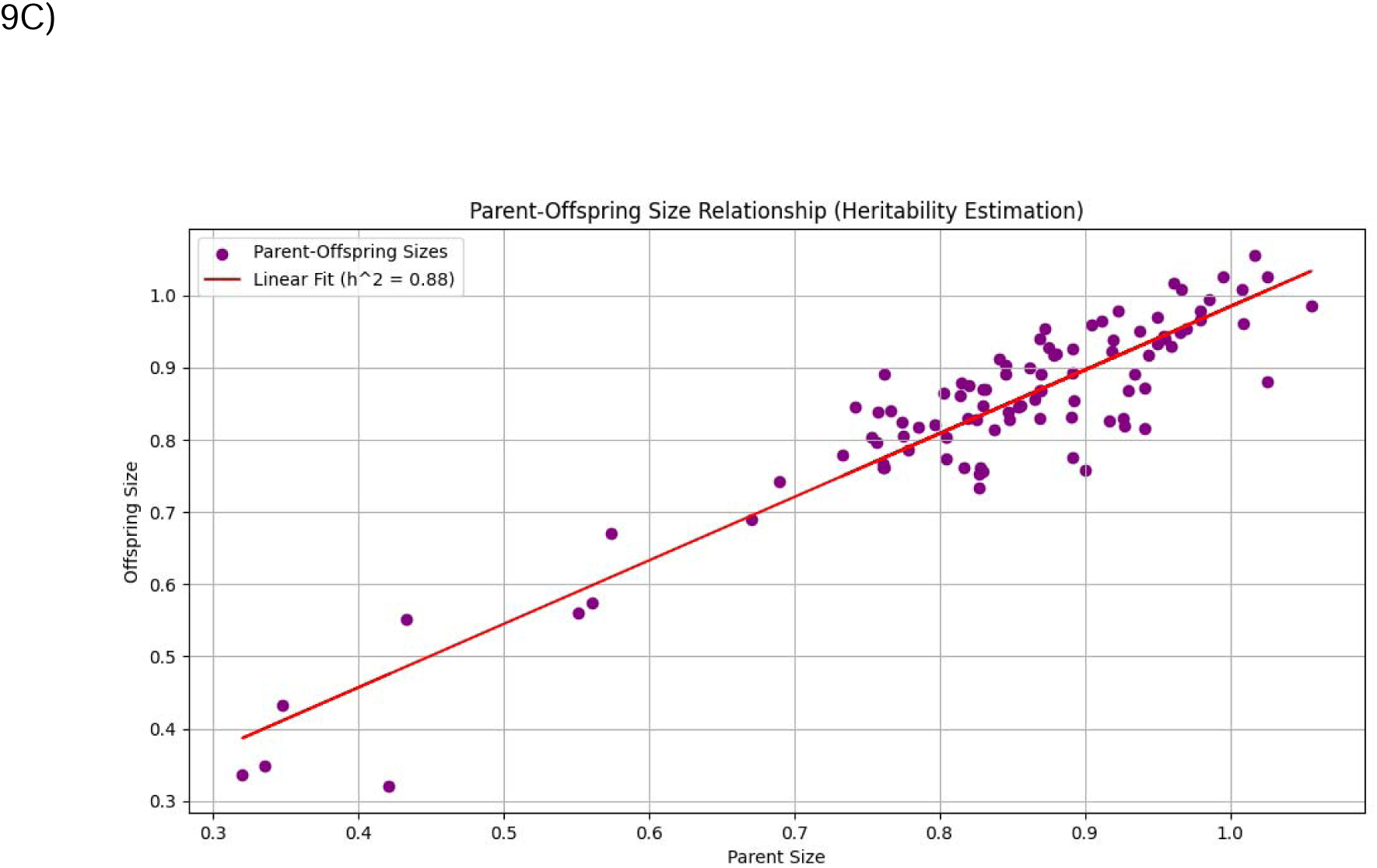
A) Continuously varying parasitoid wasp size (red), true host state (“Large” versus “Small”), and Viterbi algorithm inferred wasp host using a Hidden Markov Model (HMM) simulation, simulated over 100 generations. B) Sinusoidally varying host abundance (blue), offspring parasitoid wasp fitness (green), and efficiency of parasitization (purple) in the HMM simulation over 100 generations, and C) narrow sense heritability (h^2^ = 0.88) of body size, measured as the slope of parent-offspring linear regression in the HMM simulation over 100 generations.

**Figure 10:**
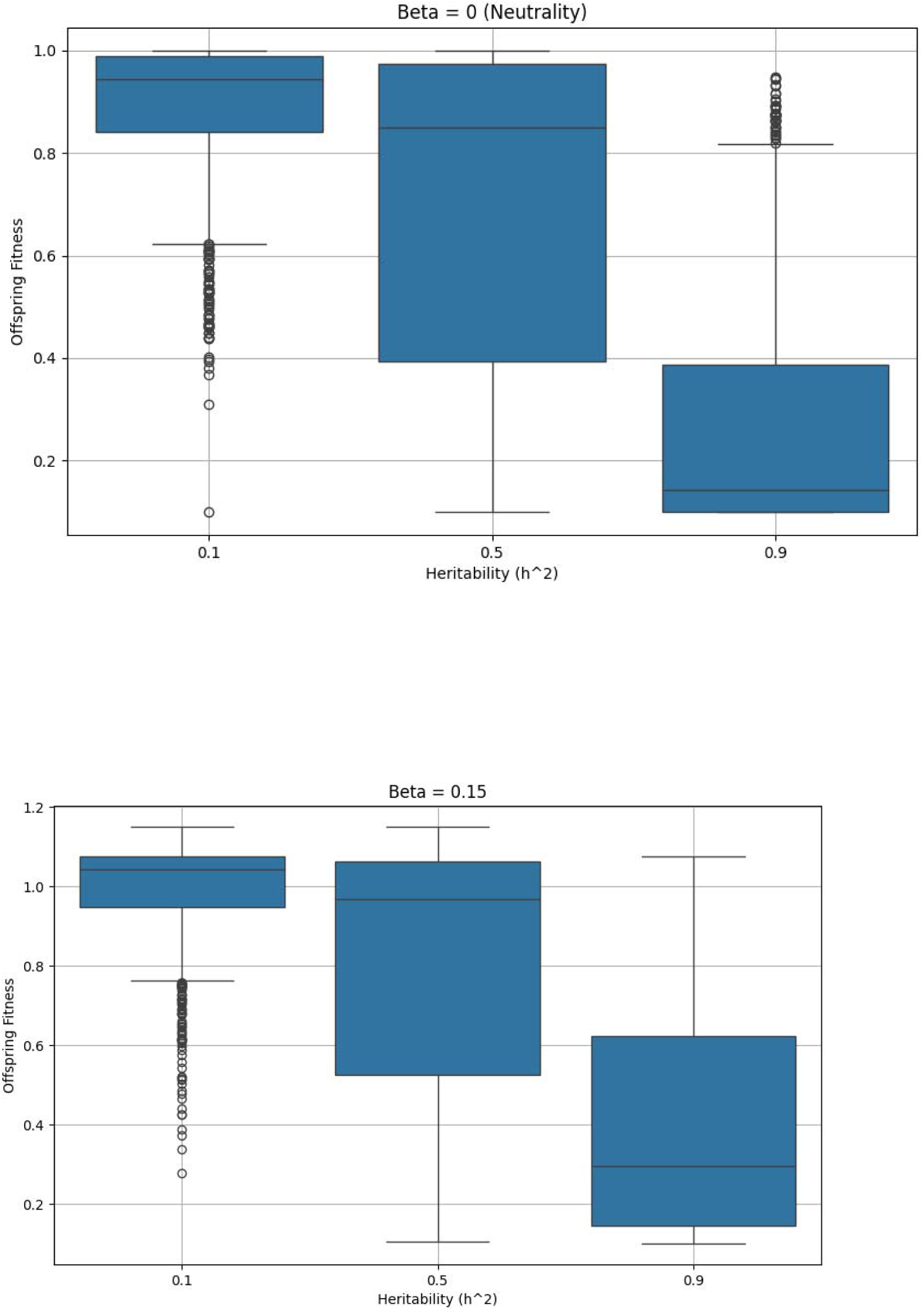
Offspring fitness measured as a function of varying heritability of size (h^2^ = 0.1, 0.5, and 0.9) under (A) neutrality (β = 0.0) versus (B) fecundity constraint (β = 0.15).

## Discussion

The unique life history strategies of *D. coccinellae* - thelytokous parthenogenesis, solitary behavior, and the ability to successfully oviposit in an uncharacteristically large range of host lady beetle species that span a wide spectrum of body sizes and shapes (Balduf, 1926; Ceryngier et al., 2012, 2018; Wright, 1979) - present a great opportunity to understand the dynamics of phenotypic microevolution of size. This parasitoid attacks a group of predatory beetles that are widely used in biological control; our study highlights the importance of examining the genetic bases of ecological interactions underlying parasitoid-host relationships (Fei et al 2023, Rodrigues et al 2022, Sentis et al 2022).

Specifically, the diversity in host coccinellid morphology offers *D. coccinellae* (1) different host-parasitoid conflicts (Orr et al., 1992), (2) different environmental niches for their larvae to develop in, and (3) varying amounts of adipose tissue to feed upon. Therefore, we would predict that phenotypic plasticity in *D. coccinellae*’s ability to successfully parasitize its hosts offers the species a selective advantage at microevolutionary scales, while an occasional sexual reproductive cycle with a male (Shaw et al., 1999) offers an “escape” from Muller’s ratchet (i.e. irreversible accumulation of deleterious variants towards extinction). It has been well documented that variation in parasitoid wasp morphology is strongly associated with variation in body size and morphology of host species (Belshaw *et al.,* 2003; Symonds and Elgar, 2013).

Furthermore, previous research indicates that the environmental variation in host lady beetle body size strongly influences the body size phenotype of each emergent *D. coccinellae*, with each next clonal generation being capable of significant size changes relative to the parent (Vansant *et al*., 2019). In this study, we utilize a common-garden, reciprocal transplant experiment over multiple generations to investigate the variation in body size morphology of emergent *D. coccinellae* conditioned on (1) the same host species (unilineal), and (2) alternating host species (multilineal). Our study clearly points to the dependence of body size morphology and phenotypes of emergent *D. coccinellae* on its clonal parent, further bolstering the idea of a plastic response to maintain size variation in the species at microevolutionary scales. As *D. coccinellae* reproduces through thelytoky, the process of asexual reproduction in which diploid female parasitoids are born from unfertilized eggs, it can reasonably be expected that body size morphometric traits would exhibit strong correlation as estimated using parent-offspring regressions, as there is no source of additional genetic variation to affect relatedness through sexual reproduction and recombination or dominance (Slobodchikoff and Daly, 1971; Heimpel and De Boer, 2008). Yet, anything but a strong relationship is observed in our results. Across both the unilineal and multilineal parent-offspring regressions, most of the relationships return non-significant linear slopes, which imply that there is no difference from regression slopes of zero, indicating that there is extremely low heritable variation of size (power analysis estimated that we had an 80% chance of detecting slopes of 0.41 or larger in parent/offspring regressions). This was an interesting finding, as we considered thelytokous parthenogenesis to be such a strong constrictor on genetic variation, that the significant shift in body size would have been expected to be at least partially evident in body size plasticity. This experiment also points to how low heritability could emerge from intense selection (here artificial). It is possible that an adult *D. coccinellae* can feasibly jump to a different species of host ladybeetle than that of their mother, given the available distribution of phenotypic variation in body size across one generation. Yet, repeated host shifts in a rapid succession of a few generations may introduce too intense an artificial selection pressure for this trait plasticity to endure, limiting the variation in body size variation of the following *D. coccinellae* generations. Therefore, as a result of negligible additive genetic variance in body size morphometric traits, we would also predict that there is little trait variability for natural selection to act on/work with, thereby minimizing the trait’s ability to evolve. This is further complemented by the lack of significant differences in body size morphometric traits in emergent *D. coccinellae* among all host types as observed in our experiment. Further analyses of our controlled reciprocal transplant experiments, to quantify the fecundity of *D. coccinellae* females, perhaps differentially across different hosts would help predict the fitness consequences of natural selection on plastic size in these predominantly asexual species.

Additionally, of potential interest then is the differential efficacy of parasitization of small *D. coccinellae* on smaller versus larger coccinellid hosts. It has been predicted that host manipulation via “bodyguard” behavior (Maure et al., 2011, Maure et al., 2013) to protect *D. coccinellae* pupae from predators is presumably under selection for larger hosts, to possibly repel larger predatory species, e.g., crickets or carabid beetles. This hypothesis can also be tested by studying the fecundity, survival, duration of “bodyguard” behavior, and parasitization rates of emergent *D. coccinellae* across different Coccinellid hosts, while controlling for host size. It has also been noted that the sex of the coccinellid host, and prey availability in the field could also influence variability in size of adults (Belnavis, 1988), which were not controlled in our study.

Multivariate comparison of mother and daughter morphology yielded evidence that mothers produce offspring that differ from them, independent of the host species. Mother’s loadings on the first canonical correlation axis were positive, while their daughter’s morphology had negative loadings, except for their wing lengths. This suggests that across all microevolutionary scenarios in our experiment, large mothers produce small daughters with long wings, while small mothers produce large daughters with short wings. A small body size with long wings is consistent with better dispersal ability (summarized in Johannson et al., 2009), and it is possible that large females are preferentially producing daughters that will disperse greater distances. Smaller mothers that produce large daughters with short wings may be maximizing the survival probability of their daughters at the expense of their potential dispersal distances.

Any advantage in dispersal ability from increased body size can be critically beneficial to adult *D. coccinellae* when the population density of their host coccinelids are low or unpredictable, as increased dispersal distance offers parasitoids more opportunity to encounter a suitable host to oviposit within (Wang and Messing, 2004). Conversely, reduced dispersal ability from body size can be beneficial in an environment rich in host population density (Wang and Messing, 2004), and the generalism of *D. coccinellae* in host species selection aids in maximizing each encounter with a host coccinellid as an opportunity to oviposit within.

This pattern of increase in size of koinobionts such as Braconid wasps has also been previously reported to be correlated with increased longevity and fecundity (Boivin 2010). For example, *Trissolcus mitsukurii* (Hymenoptera: Scelionidae) which were reared on larger hosts had higher fecundity than individual parasitoids which were reared on smaller hosts (Arakawa et al., 2004). Moreover, it was observed that the number of eggs a female *Spathius agrili* (Hymenoptera: Braconidae) oviposits is positively correlated with host size, in addition to other explanatory factors (Wang et al., 2008). Since this pattern is independent of both mother and daughter host species, it is likely that it is mediated by the mother’s state. Our observations therefore offer partial support for Darwin’s fecundity-advantage model, which posits that most female species of larger body size have higher fecundity, but limited by energy availability from the host environment (Shine 1988). The mechanism for producing these changes is unknown, but facultative changes in the size of eggs, modified fecundity on energy availability, or epigenetic regulation of gene expression are some combination of the above are possible. The phenotypic variance caused by mothers producing daughters who are genetically identical but are morphologically different would further reduce the narrow sense heritability of traits, and could explain negative slope estimates for some of the traits.

An RDA was performed on the combined unilinear and multilinear morphometric datasets with the constricted set of explanatory predictor variables. The predictor variables identified with the most explanatory power were the offspring host species *C. septempunctata* and *H. convergens*, as well as the parent wasp head depth and thorax depth morphometrics. Evaluation of the RDA model found that this combination of predictors explained approximately 15.84% of the variance in the offspring morphology response matrix. The explanatory power of the *C. septempunctata* and *H. convergens* offspring host species in explaining offspring morphometrics of *D. coccinellae* supports that the species of host coccinellid (Orr et al., 1992; Arakawa et al., 2004; Wang and Messing, 2004), in addition to host body size (Vansant, et al., 2019), is a significant factor in producing larger body size of offspring *D.coccinellae*. The significance of thorax-depth morphometrics in parent *D. coccinellae* could be related to the thorax body segment housing movement appendages, specifically the legs and wings located on the thorax along with the internal muscle tissue required to enable movement (Dudley, 2002; Fischbein, et al., 2018), with larger wasps having the advantage in dispersal ability (Ellers et al., 1998). As *D. coccinellae* allocates more energy to a larger thorax depth during developmental stages, more muscle tissue can be contained within the thorax and can increase the stored elastic energy within the thorax for movement and flight (Dudley, 2002; Fischbein, et al., 2018). Conversely, if developmental energy is not invested in increasing the thorax size, it can be invested in other body segments.

In addition, the multivariate multiple regression performed found no significant relationship between parent and offspring *D. coccinellae* morphometrics. As hypothesized, these non-significant results between parent-offspring morphometrics clearly shows no heritability of *D. coccinellae* body size, regardless of her thelytokous parthenogenetic method of asexual, clonal reproduction. Our findings from the multivariate multiple regression performed are similar to the conclusions in Bennett, D. and Hoffman, A., 1988, in which they found no evidence for the heritability of body size from their regression analysis of mother-daughter *Trichogramma carverae* (Hymenoptera: Trichogrammatidae).

We replicate these results using simulations under a Hidden Markov Model of constrained fecundity under Cope’s Law (i.e. a selection gradient for larger size), wherein larger body size, combined with the availability of a larger host offers considerable fitness advantages to wasp offspring. Additionally, plasticity of size (and ergo low heritability) contingent on host availability and size provides considerable offspring fitness advantages, even if the selection gradient (β) is 0 (neutral), or 0.15 (median selection gradient for size estimated across animals).

Finally, our results also bring into question the micro- and macroevolutionary consequences of the evolution and maintenance of thelytokous parthenogenesis from ancestral arrhenotoky in these species. A recent study on the *D. coccinellae* genome by Sethuraman et al., 2022 pointed to an early divergence, accelerated rates of genome evolution via manifold duplications and gene loss along the *D. coccinellae* lineage. Significant duplication events were reported in transposase activity and stress response gene families, while significant gene losses were reported among olfactory/odorant receptors and viral-coevolution genes. We surmise that these duplication (and loss) events contribute to standing genomic variation in *D. coccinellae* that permit plasticity of size despite parthenogenetic reproduction and alternating reproductive trade-offs depending on host availability and host-associated energy limitations, independent of maternal genetics.

**Table 1:** Parameters and variables modeled in our simulation of the evolution of body size as a function of host abundance, parasitization efficacy, host size, and fecundity constraint on size based on Cope’s Law.

| Simulation Parameter/Variable | Values/Ranges |
| --- | --- |
| Host size | Binary: Small (0) or Large (1) |
| Wasp size | Continuous: (0.0, 1.0) |
| Offspring wasp fitness | Efficacy of parasitization + Selection term;<br>Continuous (0.1, 1.0); minimum fitness = 0.1 |
| Host transition probabilities | [0.8, 0.2], [0.3, 0.7] |
| Selection gradient ($\beta$) | Continuous: (0.0, 1.0); 0 = Neutrality, > 0.0 indicating selection for larger hosts |
| Host abundance | Continuous, $1 + 0.5 * \sin(\text{generation}/10)$;<br>fluctuates in every generation between 0.5 and 1.5 |
| Optimal size of wasp | 1 if host size = 1, else 0.5 |
| Size mismatch penalty | $\exp(-((\text{wasp\_size} - \text{optimal\_size})^2) / (2 * 0.1^2))$ |
| Efficacy of parasitization | Size mismatch penalty * Host abundance |
| Selection term | Selection gradient * wasp size |
| Number of generations | (100, 1000) |
| Initial state | Random; $\sim U(0, 1)$ |
| Initial parasitoid size | 0.5 |
| Narrow sense heritability of offspring size | (0.1, 0.3, 0.9) |

## Data Availability

All high definition images utilized in assessment of size in this study, and Python code used in simulations have been made available via FigShare (https://doi.org/10.6084/m9.figshare.21685181.v1). All statistical analyses and morphological data used in this study are accessible via R Markdown at: https://wkristan.github.io/toval_etal_supplement.zip.

## Acknowledgments

This work was funded by the National Institute of Food and Agriculture, U.S. Department of Agriculture, Hatch Program under accession number 1008480 and funds from the University of Kentucky Bobby C. Pass Research Professorship to JJO, NSF ABI Development #1664918, NSF CAREER #2042516 to AS, USDA NIFA #2017-06423 to PI George Vourlitis and co-PI AS, NSF REU #1852189 to PI Betsy Read and co-PI AS, CSUSM GPSM #86969 to AS, and a CSUPERB COVID-19 Research Recovery Microgrant to AS. AS would like to thank Alicia Tovar and her mother for their selfless and invaluable help to keep this study going despite the onset of the COVID-19 pandemic - this work would have not been possible without their help. SM was supported by the NIH MARC Program at SDSU and AK was supported by startup funds to AS at SDSU and NIH1R15GM143700-01 to PI Kimberly Ayers and AS. TM was supported by a CSUBIOTECH grant to AS. This work is dedicated in the memory of Roxane Saisho who we tragically lost in December 2022 - we will forever be thankful and indebted to her contributions to this study and our lab.

